# Marmoset monkeys (*Callithrix jacchus*) develop eosinophilic airway inflammation after house dust mite exposure

**DOI:** 10.1101/2020.02.24.962498

**Authors:** F Dahlmann, C Curths, J Wichmann, S Jimenez Delgado, E Twisterling, YS Kap, N van Driel, BA’t Hart, Y Knauf, T Becker, S Dunker, H Windt, M Bleyer, FJ Kaup, K Sewald, A Braun, S Knauf

## Abstract

**Background:** Extensive analysis of eosinophilic airway inflammation in human-relevant animal models is required to test novel, human-specific pharmaceuticals. This requires species, which show high genetic homology to humans such as non-human primates. Efficacy assessment of novel human-specific biologicals in eosinophilic airway inflammation is currently performed in the cost-intensive macaque asthma model.

**Objective:** The present study investigated whether marmoset monkeys (*Callithrix jacchus*), a small-bodied non-human primate species from the New World, develop eosinophilic airway inflammation in response to house dust mite allergen exposure (HDM, *Dermatophagoides pteronyssinus*).

**Methods:** Marmoset monkeys were sensitized against HDM by subcutaneous (s.c.) injection and subsequent intratracheal (i.t.) HDM aerosol challenges. Airway and systemic immunologic reactions were monitored and sensitivity towards glucocorticoid therapy was assessed. The pulmonary immunologic response was analyzed by repetitive bronchoalveolar lavage (BAL).

**Results:** Bronchoalveolar lavage fluid (BALF) exhibited increased levels of eosinophils, mast cells, and lymphocytes, as well as interleukin (IL)-13 after HDM challenges, compared to negative controls. The systemic immunologic response was assessed in peripheral blood mononuclear cells derived from sensitized animals, which secreted increased IL-13 and IFN-γ upon allergen stimulation in contrast to non-sensitized negative control animals. Although IgE was not detectable, HDM-specific serum IgG was elevated in sensitized animals. Both airway and systemic responses were reduced by treatment with glucocorticoids. However, lung function and pathological analyses did not reveal significant differences between groups.

**Conclusion:** In conclusion, marmoset monkeys developed a mild HDM-induced eosinophilic airway inflammation useful for efficacy testing of novel human-specific biologicals.

## INTRODUCTION

About 235 million patients worldwide suffer from eosinophilic allergic inflammation like asthma and require therapeutic interventions [1]. Current therapeutics comprise glucocorticoids and long-acting beta-agonists, which in most cases efficiently control mild and moderate asthma. Severe asthma, however, is frequently therapy-resistant, and only a few monoclonal antibodies such as omalizumab, mepolizumab, and reslizumab have been approved for treatment [2]. The preclinical development of monoclonal antibodies and other human-specific biologicals directed against a particular target requires high homology in rodents. If cross-reactivity is not given, non-human primates (NHPs) are often the species of choice.

To analyze the efficacy of human-specific therapeutics in NHP models of asthma, symptoms and pathology have to coincide with human asthma patients. In patients, asthma is characterized by localized airway reactions, an increase of inflammatory markers, and airway obstruction as a result of pathological changes [3]. Sensitization is triggered by inhaled allergens like house dust mite allergen (HDM), leading to an influx of inflammatory cells into the airways. Inflammatory cells, which typically increase in airways of asthmatic patients, include eosinophils and lymphocytes (reviewed in [4]), accompanied by the T-helper 2 (Th2) cytokines IL-4, IL-5, and IL-13. Systemic markers of asthma, which can be detected in blood, include allergen-specific IgE and Th2-derived interleukins (reviewed in [5]). Airway obstruction is determined by lung function analysis and characterized by an early airway response (EAR) immediately after allergen exposure and increased airway hyperresponsiveness (AHR) towards unspecific stimuli such as methacholine [6]. As an alternative to *in vivo* lung function, these changes can also be observed in *ex vivo* precision-cut lung slices (PCLS) [7]. Underlying inflammation-induced chronic pathological changes are characterized by goblet cell metaplasia, subepithelial fibrosis, smooth muscle hypertrophy, and angiogenesis [8].

Among NHPs, asthma has extensively been investigated in cynomolgus (*Macaca fascicularis*) and rhesus macaques (*Macaca mulatta*). Both macaque species develop eosinophilic airway diseases after sensitization and repetitive challenge with HDM [9–12]. Similarly, HDM-specific IgE is detected in serum [13, 14] and systemic markers of Th2-mediated inflammation are released from peripheral blood mononuclear cells (PBMC) after stimulation [10, 11]. Macaques show HDM-mediated airway obstruction [9–11] and an increased AHR in lung function analyses after methacholine (MCh) exposure [10, 11, 15], which can also be observed in *ex vivo* PCLS [16]. Moreover, pathological changes include goblet cell hyperplasia, epithelial hypertrophy, thickening of the basement membrane, and eosinophilic infiltrations [9, 10, 17], comparable to human patients.

Other NHPs that have been investigated in inflammatory airway disease research are marmoset monkeys (*Callithrix jacchus*) (reviewed in [18]), which are smaller in body size and therefore offer a cost-efficient alternative to macaques. Marmoset monkeys developed AHR and neutrophilic airway inflammation after intratracheal challenge with lipopolysaccharide, indicating the general suitability of this species in airway disease research. Moreover, the neutrophilic inflammation was sensitive towards pretreatment with glucocorticoids and a phosphodiesterase-4 inhibitor, both *in vivo* and *ex vivo* [19, 20]. Marmosets are used, when macaques show less cross-reactivity towards a specific target. This was, for example, the case for the monoclonal antibody canakinumab [21, 22].

We hypothesize that the sensitization of marmosets with HDM leads to eosinophilic airway disease similar to human asthma, which can be used for efficacy assessment of monoclonal antibodies against human asthma. Herein we report the induction of respiratory eosinophilic inflammation in marmoset after sensitization with HDM.

## METHODS

### Animals and anesthesia

Care and housing of marmoset monkeys at the German Primate Center were in accordance with the European and national directives for animal protection (2010/63/EU, §7-9/TierSchG/7833-3). The study was approved under reference number “AZ 33.9-42502-04-14/1421” by the German Lower Saxony Federal State Office for Consumer Protection and Food Safety and had the institutional reference number “Int. 1/14” at the German Primate Center. Twenty-one male and six female marmoset monkeys were preselected by bronchoscopy and were grouped according to Table 1. Mean animal weight was 409.0±7.5 g, with an average age of 4.6±0.2 years (mean±SEM). For procedures requiring anesthesia, 0.3 mg/kg body weight (BW) diazepam (Diazepam-ratiopharm, Ratiopharm) and 10 mg/kg BW alfaxalone (Alfaxan, Vétoquinol) were injected intramuscular (i.m.). During pulmonary function testing, anesthesia was maintained with isoflurane (Baxter). Information regarding care is provided in the supplementary methods. Good Veterinary Practice was applied whenever animals were handled.

**Table 1:**
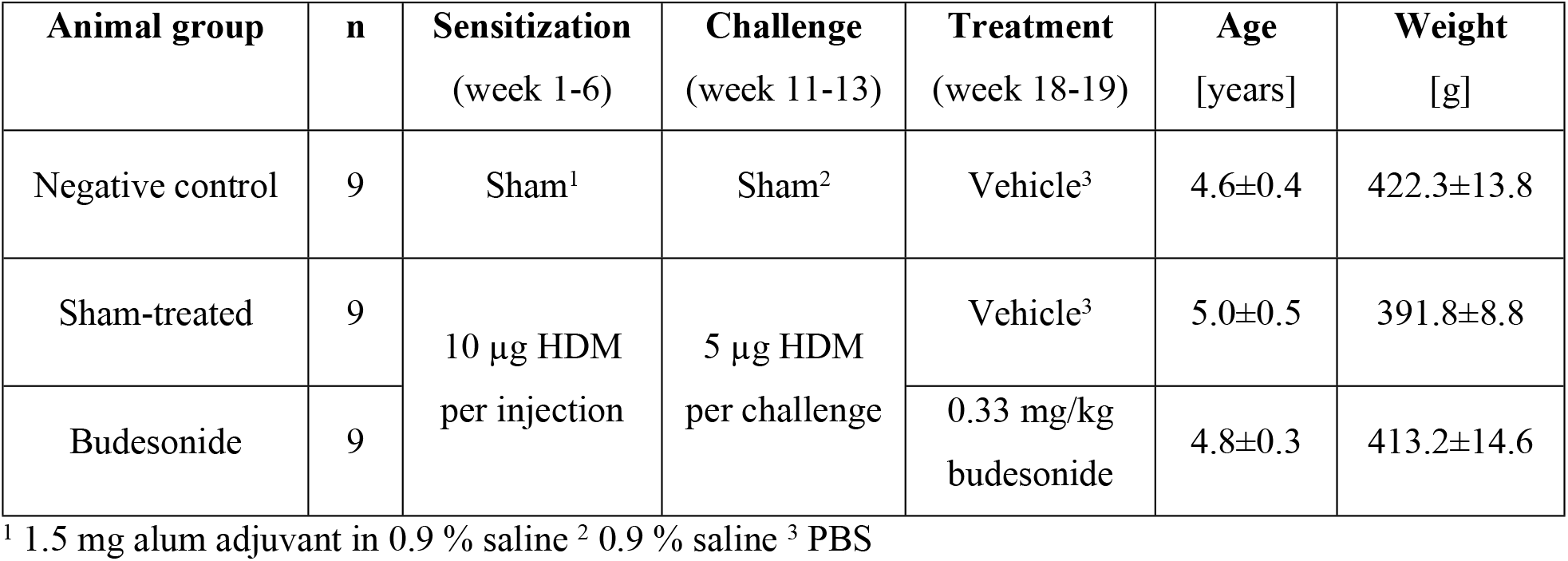
Animals used in the study. Animals were separated into three different groups (negative control, sham-treated, budesonide) according to age and body weight (mean values +/− SEM) at the beginning of the study (stratified randomization). Each group consisted of seven male and two female animals. House dust mite allergen (HDM) doses for sensitization and challenge were 10 and 5 µg, respectively. Budesonide-therapy was performed with 0.33 mg/kg body weight. For more details, see Table 2.

### Study Design

Animals were allowed to acclimatize to the experimental housing conditions for at least two weeks. Eighteen healthy marmosets were sensitized against HDM six times once a week by subcutaneous (s.c.) injection of 10 µg *Dermatophagoides pteronyssinus* crude mite extract (Greer) with 1.5 mg adjuvant (Imject™ Alum, Thermo Scientific^TM^) (Table 1, Table 2). HDM extract contained 7.3 endotoxin units per µg of protein. Negative control animals received only adjuvant (n=9). Before and after sensitization, skin prick tests were performed as described in the supplementary material. Four weeks after sensitization an aerosol challenge phase followed with six intratracheal (i.t.) applications of 5 µg allergen over three weeks. For i.t. administration, a MicroSprayer (Penn Century) was inserted into a custom-made orotracheal tube as described elsewhere [19]. The last challenge was performed only in a subset of animals (n=17), utilizing a lung function measurement device [20]. All remaining animals received HDM via the MicroSprayer. At this time point, even the negative control animals received allergen. At the end of the study, four weeks after the last challenge, the therapeutic intervention was performed i.t. on three consecutive days with either sham substance (PBS, n=17) or budesonide (0.33 mg/kg, acis Arzneimittel, n=9), followed by a final HDM challenge (5 µg) in all animals. After end-point lung function testing and subsequent bronchoalveolar lavage (BAL), animals were humanely euthanized to perform *ex vivo* and pathological analyses.

**Table 2:**
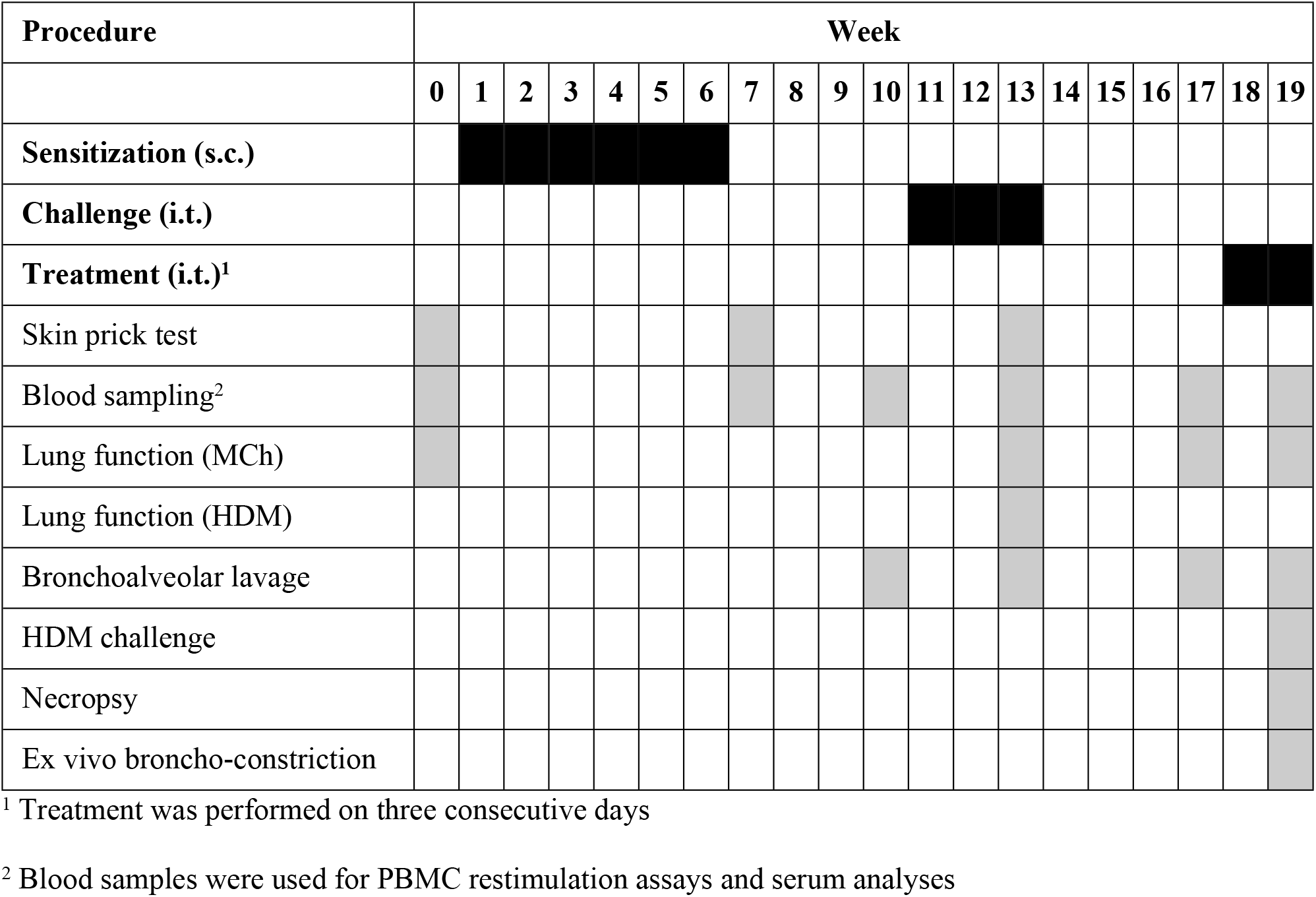
Study design. The three phases of the study are indicated as black bars. All animals underwent the described procedures at the respective time points (grey boxes), except for skin prick testing and lung function after house dust mite (HDM) exposure in week 13. Subcutaneous (s.c.) sensitization was performed once per week, whereas intratracheal (i.t.) challenge was performed twice weekly. Animals were treated i.t. according to Table 1A on three consecutive days. MCh=Methacholine

### Bronchoalveolar lavage

Video-assisted BAL was performed in anesthetized marmosets as described previously [20]. Briefly, a laryngoscope (HEINE Standard F.O. Classic+ Miller 00, Heine) was used to insert the custom-made bronchoscope (Karlheinz Hinze Optoengineering) into one of the main bronchi. The right (pre- and post-challenge) or the left (pre- and post-therapy) lung was flushed twice with pre-warmed 3 ml sterile saline solution (0.9 % NaCl, WDT). Protease inhibitor cocktail (Sigma-Aldrich) was immediately added to the recovered BAL fluid (BALF), which was centrifuged for 5 min at 340 ×*g* at 4 °C. BALF supernatants were stored at −80°C until further processing. BALF pellets were resuspended in phosphate-buffered saline (PBS) and centrifuged by a Shandon Cytospin (Thermo Scientific™) to generate thin-layer cell preparations (“cytospots”). Cytospots were stained by May-Gruenwald Giemsa stain and mast cells were detected immunohistochemically (monoclonal mouse anti-human mast cell tryptase, Dako). Staining protocols and further details can be found in the supplementary material. Total and differential cell counts were evaluated by microscopic counting of 200 cells per sample.

### Blood sampling, PBMC isolation, and restimulation

In conscious animals, blood was collected from the *Vena femoralis* and immediately transferred into EDTA (Vacutainer Glass K_3_ EDTA tubes, BD) or serum tubes (Vacuette Serum Clot Activator tubes, Greiner Bio-One), respectively. EDTA blood was processed for PBMC isolation and was additionally analyzed in an automated hematology system (Advia 2120, Siemens). Serum was centrifuged (15 min, 2000 ×*g*, room temperature) and stored at −20 °C for immunoglobuline and C-reactive protein (CRP) quantification (Dimension Xpand Plus, Siemens).

Whole blood was diluted with PBS (PAN^TM^ Biotech) and loaded on Lymphoprep™ medium (Stemcell^TM^ Technologies). Tubes were centrifuged (20 min, 800 ×*g*, room temperature) and the mononuclear cell layer was transferred to PBS and washed twice. PBMC pellets were resuspended in cell culture medium (RPMI 1640, PAN^TM^ Biotech) that contained 10 % fetal bovine serum (FBS, Biochrom AG) and 1 % Penicillin/Streptomycin (PAN^TM^ Biotech). PBMC were incubated overnight in 96-well microplates (Cellstar, Greiner Bio-One) with 2×10^5^ cells per well at 37 °C and 5% CO_2_. Thereafter, PBMCs were stimulated with 10 µg/ml HDM for 96 h. Controls involved medium or HDM plus 50 µg/ml dexamethasone (Dexa-ratiopharm, Ratiopharm GmbH). Supernatants were stored at −80 °C and were used for cytokine detection by ELISA.

### ELISA

Supernatants from BALF and PBMC restimulation assays were analyzed for IL-13 and IFN-γ levels, using commercially available marmoset-specific ELISA (Marmoset IL-13 and IFN-γ ELISA Ab pair, U-CyTech). Assays were performed according to the manufacturer’s protocols. Detection limits were set to lower limits of quantification and revealed ≤0.64 pg/ml (IL-13) and ≤3.82 pg/ml (IFN-γ) for PBMC samples and 0.32 pg/ml (IL-13) and ≤0.70 pg/ml (IFN-γ) for BAL samples.

Allergen-specific IgG was detected using a custom-made ELISA. Briefly, 96 well plates (Greiner Bio-One B.V.) were coated with 5 µg/ml HDM extract overnight and blocked with PBS supplemented with 2 % BSA at 37 °C for 1 h. Samples were diluted 1:100 and incubated for 2 h at 37 °C. A polyclonal alkaline phosphatase-conjugated rabbit-anti-human IgG (Abcam^®^), diluted 1:2000 in 1 % BSA in PBS was added for 1 h at 37 °C. Substrate p-Nitrophenyl phosphate (SIGMAFAST™, Sigma-Aldrich) was added for 20 min. Optical density (OD) was subsequently measured at 405 nm (BioTek Instruments Inc.). Washing steps were performed using 0.05 % tween 20 in PBS. Antibody levels are given as arbitrary units (AU), derived from comparison to standard curves from control marmoset serum. The Ab content in the pooled plasma was defined at 2.500 arbitrary units, and newly collected ELISA data were fitted to a four-parameter hyperbolic function using the homemade ADAMSEL program developed by Dr. E. Remarque (Biomedical Primate Research Centre, the Netherlands).

### Lung function measurement

MCh tests were conducted before sensitization, after the challenge phase, and prior and after therapy, whereas *in vivo* reaction against HDM was only assessed after challenge in a subset of animals. *In vivo* assessment of lung function was performed as previously described [20]. Briefly, anesthetized spontaneously breathing animals were orotracheally intubated and connected to the lung function device. Pulmonary parameters including lung resistance (R_L_) and dynamic compliance (C_dyn_) were recorded to assess AHR by exposure to MCh. Both parameters were described as the provocative dose (PD) of MCh and are analyzed as increased R_L_ 150% above baseline (PD_150_R_L_) or decreased C_dyn_ 50% below baseline (PD_50_C_dyn_). Reduced PD values are indicative for AHR. Animals were provoked with either increasing aerosol doses of 0.25 µg to maximal 64 µg MCh (acetyl-β-methylcholine chloride, Sigma–Aldrich) or a single application of 5 µg HDM. MCh application was stopped when R_L_ reached 150 % above the individual’s baseline. Subsequently, animals received 0.1 mg salbutamol (Ratiopharm) to smoothen recovery. For lung function analysis Notocord-hem^TM^ software (NOTOCORD Systems, Croissy Sur Seine, France) was used.

### Necropsy, histopathology, and immunohistochemistry

Euthanasia was performed in deep anesthesia using an intracardial injection of a lethal dose of pentobarbital sodium (Narcoren, Merial). Post-mortem, the thoracic cavity was opened and the thoracic content was removed en bloc. The right caudal lung lobe was removed, cannulated and instilled with 4% phosphate-buffered formaldehyde at 25-cm hydrostatic pressure. After floating fixation for at least 24 h, the right caudal lung lobe was embedded in paraffin and further processed by hematoxylin and eosin (H&E) stain and secretory cell detection by immunohistochemistry (IHC). Further details are provided in Supplementary Methods.

### Ex vivo bronchoconstriction of PCLS

PCLS were generated from five animals per group after therapy as previously described [16]. Briefly, the left lung was filled with 1.5 % low-gelling temperature agarose (Sigma-Aldrich) in cell culture medium (MEM HEPES Modification, Sigma-Aldrich) and hardened in ice-cold PBS. PCLS of 8 mm diameter with cross-sectioned airways were generated using a Krumdieck tissue slicer (Alabama Research and Development) and incubated in DMEM F12 (Gibco) including 100 units/ml penicillin, 100 µg/ml streptomycin, and 25 mM Hepes (Lonza) at 37 °C and 5 % CO_2_. The degree of bronchoconstriction towards 10 μg/ml HDM or 10^−7^-10^−3^ M MCh was monitored using video microscopy (Stereo microscope Zeiss Discovery V8, Digital video camera AxioCam IcM1 and AxioVision Discovery 4.8.2 Software, Zeiss). Percentage differences related to initial airway areas were calculated using the ImageJ 1.44p software (National Institutes of Health).

### Statistics

Statistical analyses were performed using Prism 6.0 (GraphPad Software). Data are shown as means with a standard error of the mean (SEM), whereas non-normally distributed data are depicted as box-and-whisker plots including medians. Box-and-whisker boxes represent 25th to 75th percentiles and whiskers show maximum and minimum values. Statistical significance was determined employing parametric (paired, unpaired t-test) or non-parametric tests (Wilcoxon matched-pairs signed rank test, Mann-Whitney-Wilcoxon test). Non parametric test was used when data were not Gaussian distributed. Correlation analyses employed Pearson correlation coefficients (r). We regarded P-values of ≤0.05 as statistically significant.

## RESULTS

### Respiratory inflammatory response

To analyze the respiratory inflammatory response, marmoset monkeys underwent BAL before and after the challenge as well as before and after the treatment. After HDM challenge sensitized animals showed significantly increased eosinophil (8.4 × 10^4^ cells/ml) and mast cell numbers (1.8 × 10^3^ cells/ml) compared to the negative control animals (1.4 × 10^4^ cells/ml and 2.9 × 10^1^ cells/ml, respectively) (Fig. 1A). In contrast, negative control animals revealed elevated BALF neutrophil numbers (2.1 × 10^4^ cells/ml) compared to sensitized animals (9.5 × 10^3^ cells/ml). In BALF supernatant, IL-13 was elevated in HDM sensitized (0.66 vs. 1.02 pg/ml) (Fig. 1B), but not in negative control animals. BALF IFN-γ levels were not changed between sensitized and non-sensitized animals (Fig. 1C).

**Figure 1:**
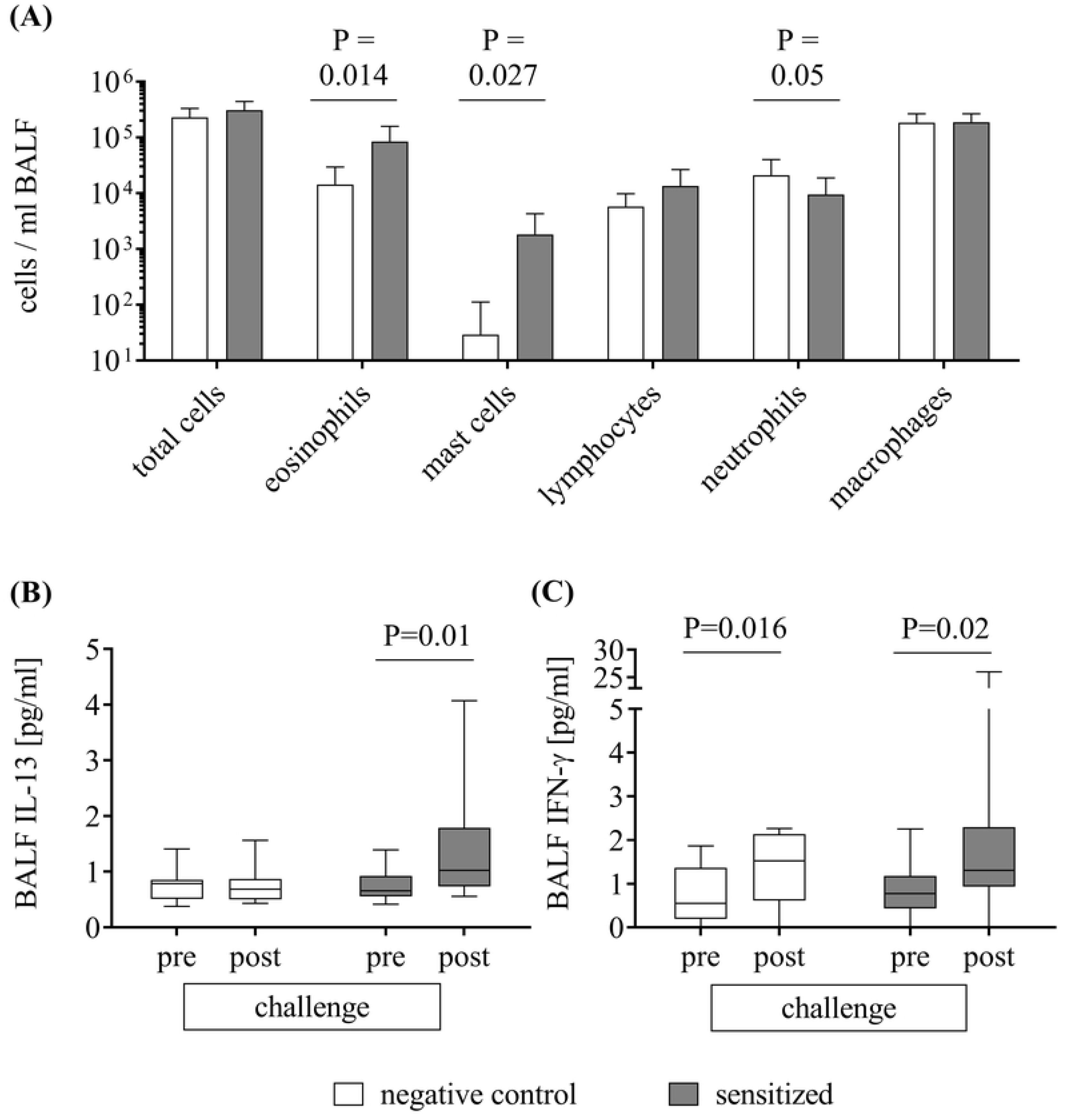
Bronchoalveolar lavage cell counts and cytokines before and after the HDM challenge phase. Bronchoalveolar lavage was performed before and after the HDM challenge phase in negative control and HDM-sensitized animals. Total cells and differentiated cell counts per ml bronchoalveolar lavage fluid (BALF) are depicted in (A). Eosinophils and mast cells were significantly increased in sensitized compared to negative control animals, whereas neutrophils were elevated in negative control animals. Cytokine analysis in BALF revealed an increase of IL-13 only in sensitized animals (B) and of IFN-γ in both groups (C). Mean+SEM; n=8 negative control, n=17 sensitized; BALF=bronchoalveolar lavage fluid; unpaired t test (A). Box plot with median±SEM; Wilcoxon matched-pairs signed rank test (B, C).

HDM-sensitized animals were treated with budesonide. In budesonide-treated animals, eosinophils were decreased compared to untreated animals (2.7 × 10^4^ cells/ml) (Fig. 2B). In contrast to the sham-treated animals, neutrophils were elevated in budesonide-treated animals (7.9 × 10^4^ cells/ml) (Fig. 2C). Additionally, a reduction of mast cells (7.3 × 10^2^ cells/ml vs. 5.4 × 10^2^ cells/ml) and lymphocytes (1 × 10^4^ cells/ml vs. 7.1 × 10^3^ cells/ml) were observed in budesonide-treated animals (Fig. 2D and E). No differences were observed for macrophages, IL-13, and IFN-γ levels before and after therapeutic intervention.

**Figure 2:**
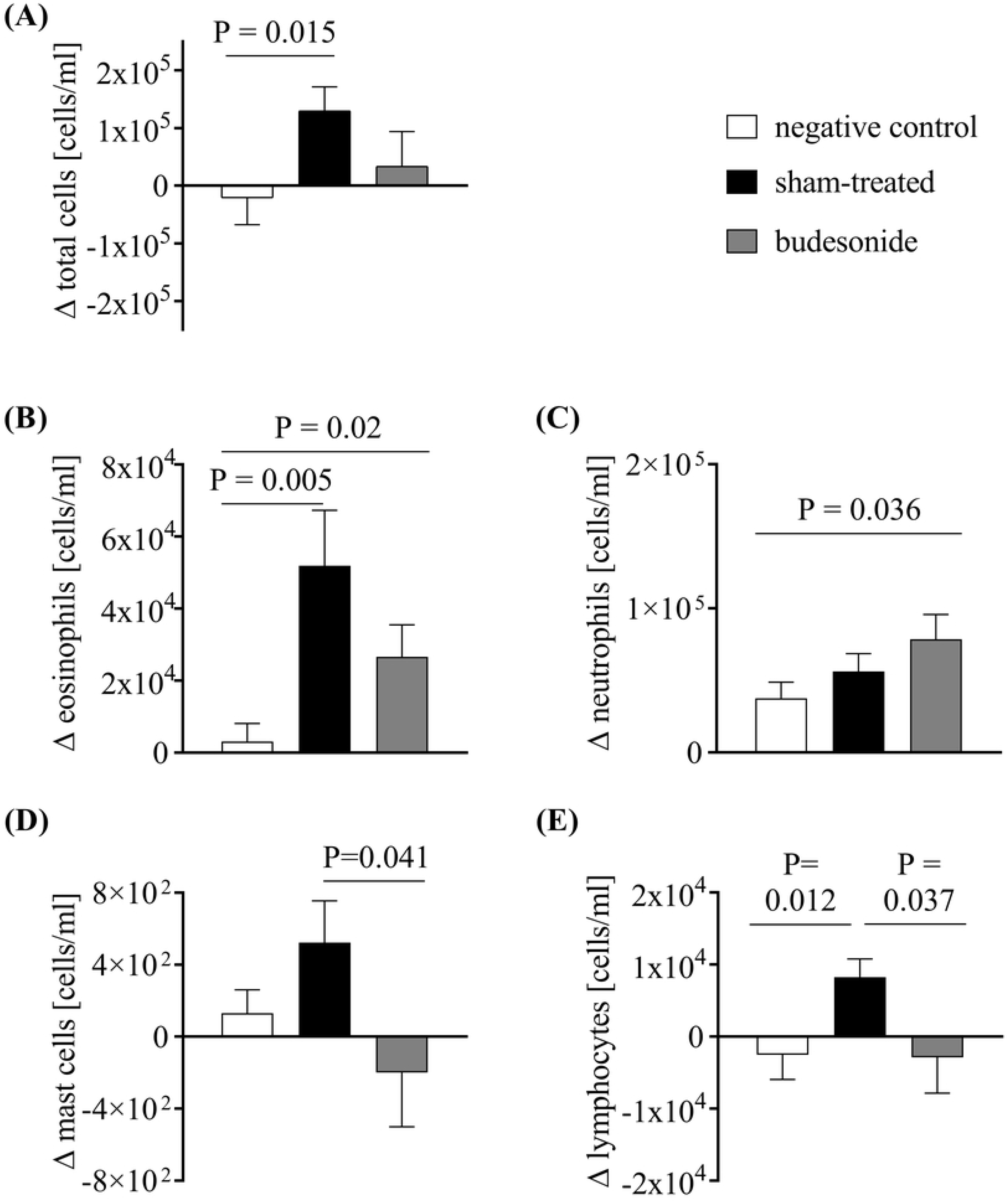
BALF cell count differences after therapeutic intervention. Bronchoalveolar lavage was performed before and after treatment and differences in cells/ml BALF were evaluated between both time points. After the treatment phase, total cell count was increased in BALF of sham-treated animals (A). Eosinophils (B), mast cells (D), and lymphocytes (E) rose to the greatest extent in sham-treated animals and were reduced by budesonide treatment. Neutrophils (C) increased in budesonide-treated animals compared to negative control animals. Mean±SEM; n=8/8/9; BALF=bronchoalveolar lavage fluid; unpaired t-test.

The allergen response towards HDM was tested by skin prick testing. Before sensitization, cutaneous reactivity tests were negative except for one out of 27 animals. Following sensitization, reactivity towards allergen was not observed in any negative control animal, whereas seven sensitized animals developed wheals against HDM (Supplementary Table S2). A correlation between positive skin prick tests and eosinophils in BAL was not observed.

Histologically, there was minimal to moderate peribronchial and peribronchiolar inflammation in the right lungs of all animals without correlation to treatment or group. The inflammatory cell infiltrate mainly consisted of lymphocytes as well as eosinophilic and neutrophilic granulocytes. There were no differences in distribution, intensity, and composition of the inflammatory reaction across all groups. An increase of tissue eosinophils or signs of hypercrinism in the sham-treated group compared to the negative control group were not noticed.

Immunohistochemical examination revealed no obvious intergroup differences in the expression of MUC5AC of the airway epithelial. In contrast, there seemed to be a decrease in the number of CCSP positive cells in the intrapulmonary airway epithelium of sham-treated animals, compared to negative control animals and the budesonide-treated group (Supplement Fig. 4).

### Systemic response: cytokines, HDM-specific IgG in serum

The systemic response was analyzed by measurement of peripheral leukocytes, immunoglobulin, and cytokine release of PBMC. Leukocyte blood cell count (4.6 x10^3^/µl vs. 6.1 x10^3^/µl) and serum CRP levels (7.6 mg/l vs. 8.6 mg/l) were significantly increased after challenge in sensitized animals (Supplement Fig. S3). Despite several attempts, total IgE and HDM-specific IgE were not detected in the blood of sensitized marmosets. However, we could show that serum HDM-specific IgG levels increased significantly after sensitization and challenge (1.7 to 5.1 AU) (Fig. 3). HDM-specific IgG was not present in the negative control animals.

**Figure 3:**
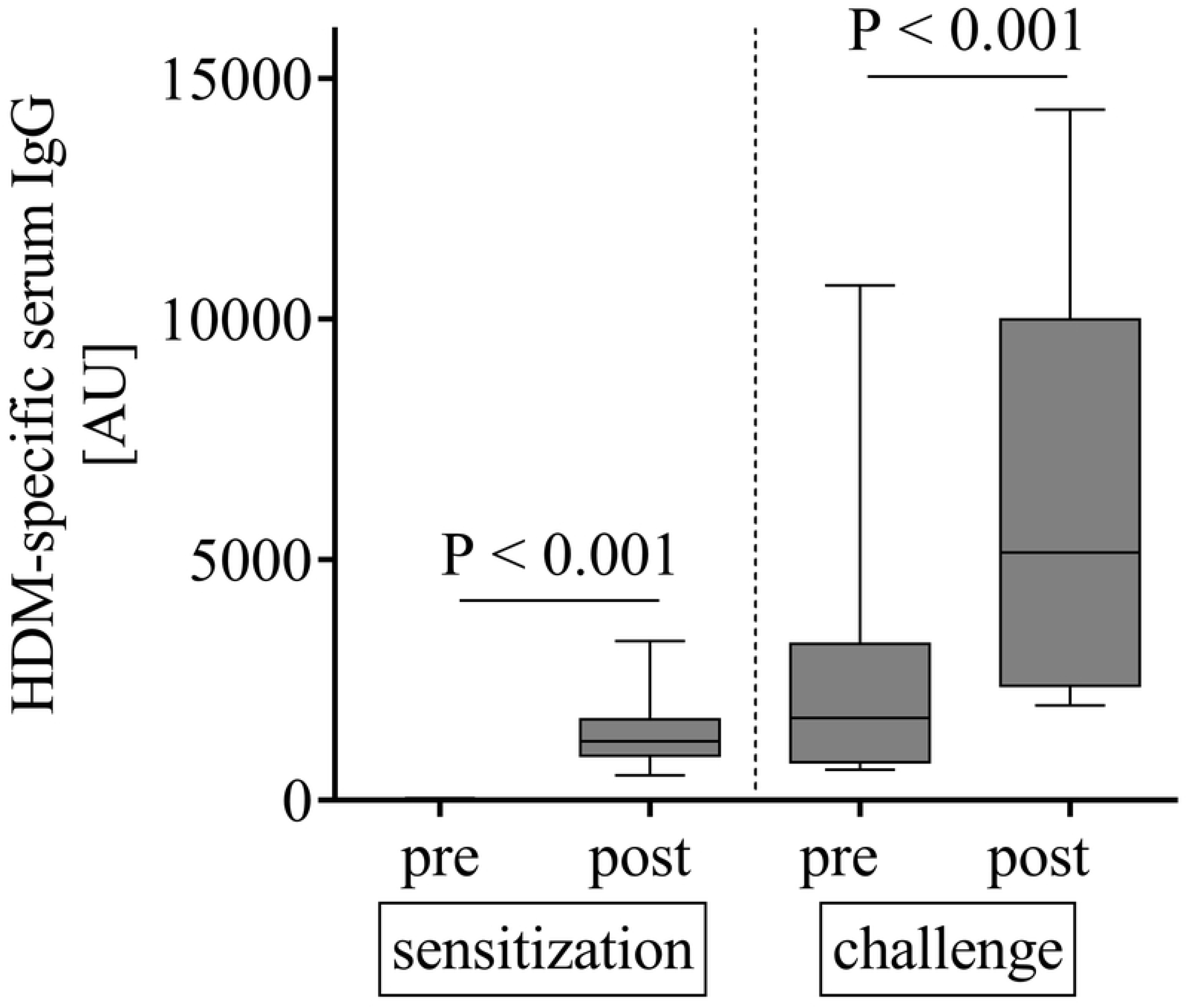
Development of anti-HDM IgG serum antibodies in marmosets. Serum IgG directed against the allergen was detected after sensitization. The HDM-specific IgG increased over time and reached a maximum after the challenge phase. Box plot with median; n=15-16; AU=arbitrary units; Wilcoxon matched-pairs signed rank test.

Cytokine release was analyzed from PBMC after *in vitro* restimulation with the allergen HDM. Whereas PBMC of negative control animals did not show a release of IL-13 or IFN-γ, PBMC of sensitized animals showed a significant increase of IL-13 (67.1 pg/ml) and IFN-γ (29.3 pg/ml) 96 h after HDM stimulation compared to medium controls (Fig. 4A and B). It was possible to attenuate these effects by adding the glucocorticoid dexamethasone to the cultures (Fig. 4A and B). Similar results were obtained after challenge and therapy (Supplementary Fig. S1).

**Figure 4:**
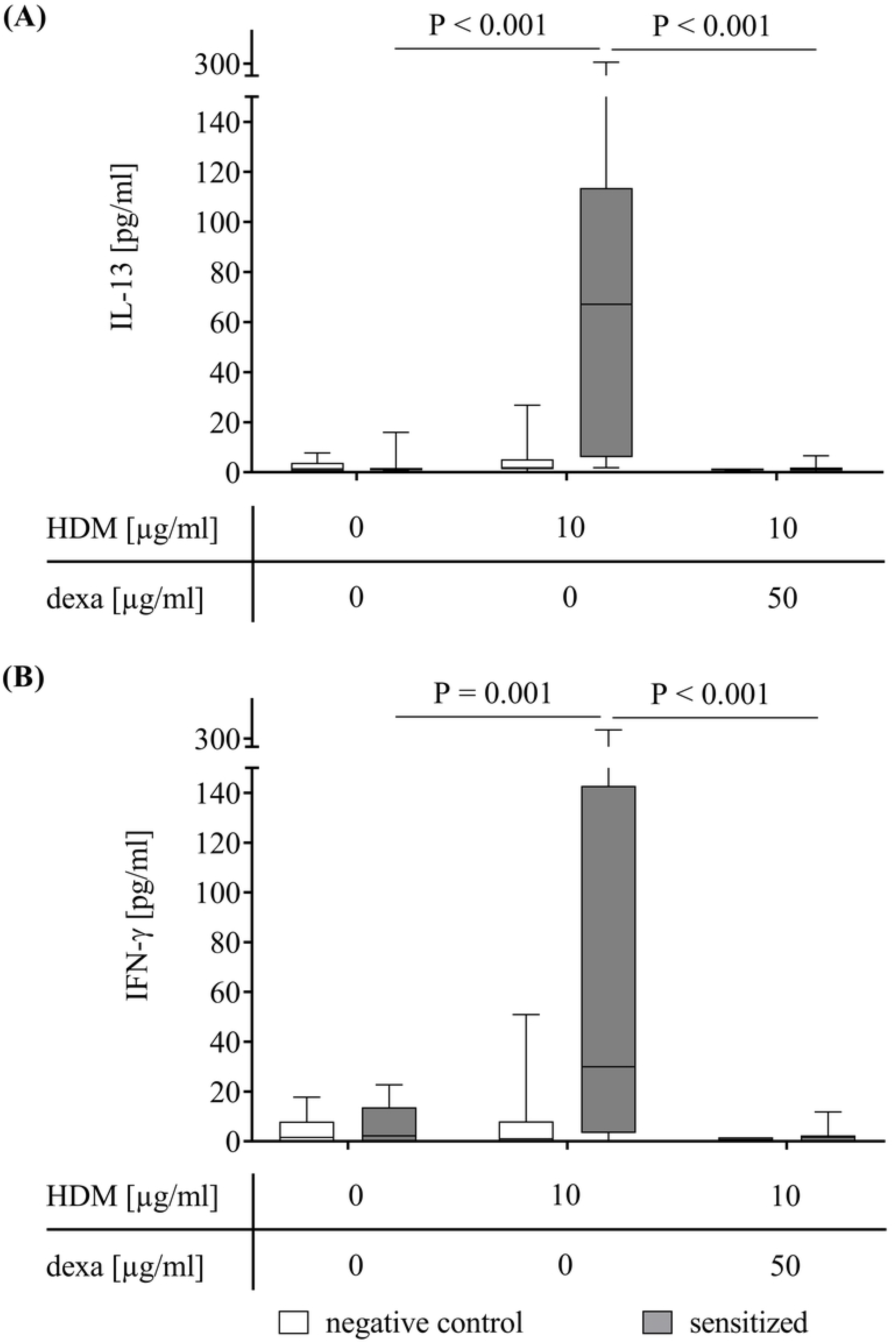
HDM-restimulated PBMC of sensitized marmosets release IL-13 and IFN-γ after allergen sensitization. PBMC were isolated after sensitization and stimulated with HDM for 96 h. PBMC from sensitized animals (grey bars) secreted elevated IL-13 (A) and IFN-γ (B) compared to unstimulated PBMC. Dexamethasone reduced allergen-related cytokine release. Allergen-dependent cytokine release from PBMC of negative animals was not elevated (white bars). Box plot with median; n varies for different stimulations; HDM=house dust mite; dexa=dexamethasone; Mann-Whitney-Wilcoxon test.

### Functional lung response

Lung function analysis included assessment of AHR by exposure to MCh at baseline, after challenge and before and after treatment. Reduced PD values are indicative of AHR. At baseline, there were no marked differences in PD_150_R_L_ between the different study groups (negative control 0.72 µg, sensitized animals 0.73 µg), which was equally the case after the challenge phase (Supplementary Table S1). Similar results were obtained for PD_50_C_dyn_. The presence of EAR towards HDM was analyzed immediately after the sixth challenge, depicted as percent change of R_L_ and C_dyn_ from baseline, but did not reveal differences. These results indicate an absence of both AHR and EAR in sensitized compared to negative control animals.

To further investigate reactivity towards the allergen, bronchoconstriction was analyzed using PCLS. Only lung tissue from sham-treated, sensitized animals showed a contraction after HDM exposure (22.7 %; Fig. 5). AHR towards MCh, as shown by EC_50_, was more pronounced in PCLS that were received from sham-treated, sensitized animals compared to negative control animals. This, however, was not significant. PCLS contraction (% airway area) upon HDM-stimulation correlated with BALF eosinophil levels (Supplement Fig. 2).

**Figure 5:**
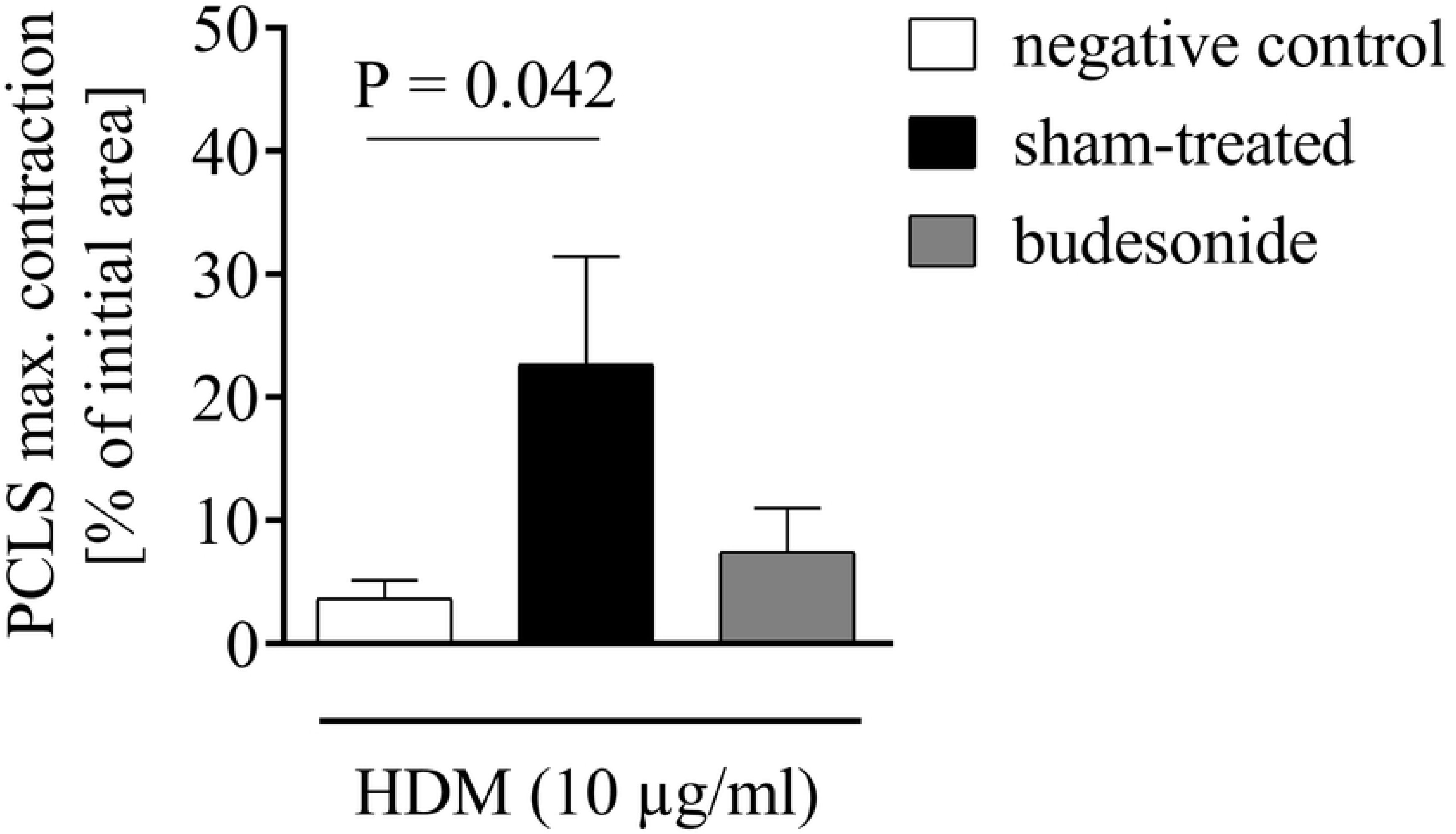
*Ex vivo* bronchoconstriction towards allergen in precision-cut lung slices. After the therapeutic intervention, PCLS were generated and stimulated with HDM. Immediately after allergen stimulation, airway areas were monitored by video microscopy to evaluate maximum contraction. Only PCLS from sham-treated animals showed a marked HDM-dependent decrease in airway area. Mean+SEM; n=5; PCLS= precision-cut lung slices; paired t-test.

## DISCUSSION

We analyzed whether marmosets develop asthma-like pathology after repeated HDM exposure and investigated sensitivity towards treatment. HDM allergen is frequently used in NHP and rodent models of asthma (reviewed in [23]) due to its relevance in human allergic asthma. We applied a systemic route of sensitization with a dose of 10 µg HDM extract, followed by intratracheal challenges with 5 µg aerosolized HDM extract in adult animals. The route was similar to other NHP models [11], although alternative exposure routes have also been reported, such as intranasal exposure [9]. The allergen challenge is required to induce respiratory inflammation [9–11, 23, 24]. The allergen amounts of HDM that were used in this study corresponded to what has been used in other NHP studies [25]. In our marmosets, the time of exposure resulted in mild airway inflammation. Most likely and based on a comparable NHP study [9], higher amounts of the allergen might have resulted in a more severe asthmatic condition.

Eosinophils, lymphocytes, and mast cell numbers are increased in the sputum and BAL of asthmatic patients [26]. Depending on the phenotype and severity of asthma, neutrophils are also increased (reviewed in [27]). Likewise, in the lungs of asthmatic marmosets, respiratory inflammation was dominated by eosinophils, mast cells, and lymphocytes in BAL. Other pathological changes such as goblet cell metaplasia, smooth muscle hypertrophy, or subepithelial fibrosis were not seen. Yet, infiltration of lung tissue with eosinophils, neutrophils, and lymphocytes was also observed in other asthmatic NHP models [9–12, 28–30].

PBMCs of asthmatic marmosets showed an increase of IL-13, IFN-γ, HDM-specific IgG, and CRP after restimulation with HDM *ex vivo*. Detection of increased IL-13 and IFN-γ in sensitized marmosets is in line with observations in human studies [31].

We were not able to detect IgE in HDM sensitized marmosets. Moreover, there is no commercial assay available for marmoset IgE and the combination of different antibodies in a self-made ELISA was not successful. As a result, we are currently not able to conclude that marmoset IgE does not play a role in our HDM sensitized marmosets. Yet, HDM sensitized marmosets showed increased levels of IgG directed against HDM. This has also been described for other NHP asthma models [10, 25] as well as increased IgG levels are reported from human asthmatics [32, 33]. We observed correlations between HDM-specific serum IgG and BALF eosinophils in sensitized marmosets (Supplementary Figure S2), which may link sensitization to respiratory inflammation. Besides the increase of antibody titers, the humoral response showed an increase of CRP in allergic animals (Supplementary Figure S3). As an acute-phase protein, CRP is generally induced by IL-6 and has been reported to be increased in allergic patients [34]. CRP is indicative of early systemic pro-inflammatory response in humans as well as marmosets.

The protocol for sensitization used in our study did not lead to impairment of *in vivo* lung function. Neither an unspecific AHR nor an EAR was observed, in contrast to what has been published in several other NHP models [9, 10, 13, 35]. However, marmosets have been shown to react with impaired lung function in a model of LPS-induced respiratory inflammation [18]. Nevertheless, EAR measured *ex vivo* by video microscopy in PCLS from allergic marmosets showed a decrease in airway area after HDM exposure. This confirms atopy in the tissue of the allergic animals. PCLS of animals pre-treated with a glucocorticoid did not show a reduction in airway area anymore, indicating an efficient therapeutic intervention.

## CONCLUSION

We report a marmoset asthma model with allergen induced mild respiratory inflammation and increased IL-13 in BAL. Compared to the old world monkey models, our marmoset model is an attractive alternative for pre-clinical drug testing of human-specific therapeutics that target the asthma-related features of the marmoset model.

## ACKNOWLEDGEMENTS

We thank K. Lampe for veterinary assistance, N. Bertram and M. Ellrodt for taking care of the animals, A. Heistermann for technical support, C. Boike, L. Erffmeier, L. Hummel, and all other members of the former Pathology Unit at the German Primate Center for technical assistance. We acknowledge the support in experiments at Fraunhofer by O. Danov, S. Konzok, H. Obernolte, S. Schindler, and E. Spies. Moreover, we thank HG. Hoymann, A. Eggel, Sophie Thiolloy, S. Wronski, E. Schelegle, K. Haanstra, M. Müller, J. Hohlfeld and R. Pabst for valuable discussion and J. Veuskens for support. We likewise thank the members of the Primomed Consortium for fruitful discussions.

## SUPPLEMENTARY METHODS

### Skin prick test

Animals were anesthetized before skin prick testing. The abdomen and the chest were clipped and disinfected, and animals were put in a dorsal recumbence. Solutions including histamine as a positive control (ALK-Abelló Arzneimittel GmbH) and HDM extract to test sensitization were dropped onto the skin, followed by a skin prick. Any developing wheal was measured in size 15 minutes after pricking the skin.

### Cytospot processing

After centrifugation, slides were air-dried and stained within the next days. Differential cell counts were analyzed after May-Gruenwald Giemsa stain. Therefore, slides were stained for 5 min with May-Gruenwald’s eosin-methylene blue solution, rinsed with ultrapure water (Co-med Online-Shop, Heusweiler, Germany) and stained for 15 min with 1:20 diluted Giemsa’s azur-eosin-methylene blue solution (Merck KGaA, Darmstadt, Germany). Slides were rinsed, air-dried and mounted using Eukitt® quick-hardening mounting medium (Sigma-Aldrich®, St. Louis, USA). Mast cells in BALF were detected immunohistochemically employing a 1:200 diluted monoclonal anti-human mast cell tryptase antibody (Dako, Hamburg, Germany). Differential cell counts were conducted on a percentage basis and cell counts per ml BALF were calculated comprising the total cell number.

### Histopathology, and immunohistochemistry

After floating fixation of the right caudal lung lobe for at least 24 h, the right caudal lung lobe was embedded in paraffin. Sections of 3-5 µm thickness were prepared from two defined localizations per animal. Specimens were stained with hematoxylin and eosin (H&E) and evaluated for the presence of inflammatory changes. Goblet cell detection was performed by immunohistochemistry (IHC) using an anti-mucin 5AC-antibody (MUC5AC, clone 45M1, Novus Biologicals; dilution 1:50). Mucin 5AC is a marker for goblet cells in human airways [36] and is also expressed in marmoset airway epithelium [37]. Also, an anti-CCSP-antibody (Clara Cell Protein Human Rabbit Polyclonal Antibody, Biovendor; dilution 1:2000) was used for evaluation of changes in airway epithelial expression of CCSP in response to allergen challenge and budesonide treatment, respectively, as previously described for humans and mice [38, 39]. IHC was performed with an automated immunostaining system (Discovery XT, Roche Diagnostics GmbH) using the SABC (Streptavidin-Biotin-Complex) method and DAB (diaminobenzidine tetrahydrochloride) for signal detection (DAB Map Kit, Roche Diagnostics GmbH).

---

**Table S1:** Marmoset *in vivo* lung function data for HDM and MCh provocations. *In vivo* lung function was evaluated at baseline (day −1) and post-challenge in negative control and sensitized animals. Provocative dose (PD) values of MCh that resulted in a 150% increase in R_L_ or a 50% decrease in C_dyn_ were assessed. At the end of the challenge phase, lung function was additionally analyzed after HDM provocation. Delta lung resistance (R_L_) and dynamic compliance (C_dyn_) percentages represent the maximum increases and decreases, respectively. Medians; n=5-9 for negative control, n=11-18 for sensitized; MCh=methacholine.

|  | baseline | post-challenge |  |
| --- | --- | --- | --- |
| $R_L$ | PD<sub>150</sub> [μg MCh] | PD<sub>150</sub> [μg MCh] | delta $R_L$ HDM [%] |
| negative control | 0.72 | 0.41 | 70.7 |
| sensitized | 0.73 | 0.49 | 24.9 |
| $C_{dyn}$ | PD<sub>50</sub> $C_{dyn}$ [μg MCh] | PD<sub>50</sub> $C_{dyn}$ [μg MCh] | delta $C_{dyn}$ HDM [%] |
| negative control | 0.37 | 0.34 | -33.1 |
| sensitized | 0.46 | 0.47 | -30.7 |

**Table S2:** Skin prick tests in marmosets. Skin prick tests were performed at baseline, post-sensitization, and post-challenge. All animals were tested before and after sensitization, whereas only two-thirds of the animals were tested post-challenge. Wheal development towards HDM extract (Greer, ALK) in respect of all valid tests are depicted. Validity was determined depending on results of control substances. Numbers indicate positive single animals of all tested animals.

| animal group | baseline | post-sensitization | post-challenge |
| --- | --- | --- | --- |
| negative control | 0/9 | 0/8 | 0/5 |
| sensitized | 1/18 | 7/17 | 4/9 |

**Figure S1:**
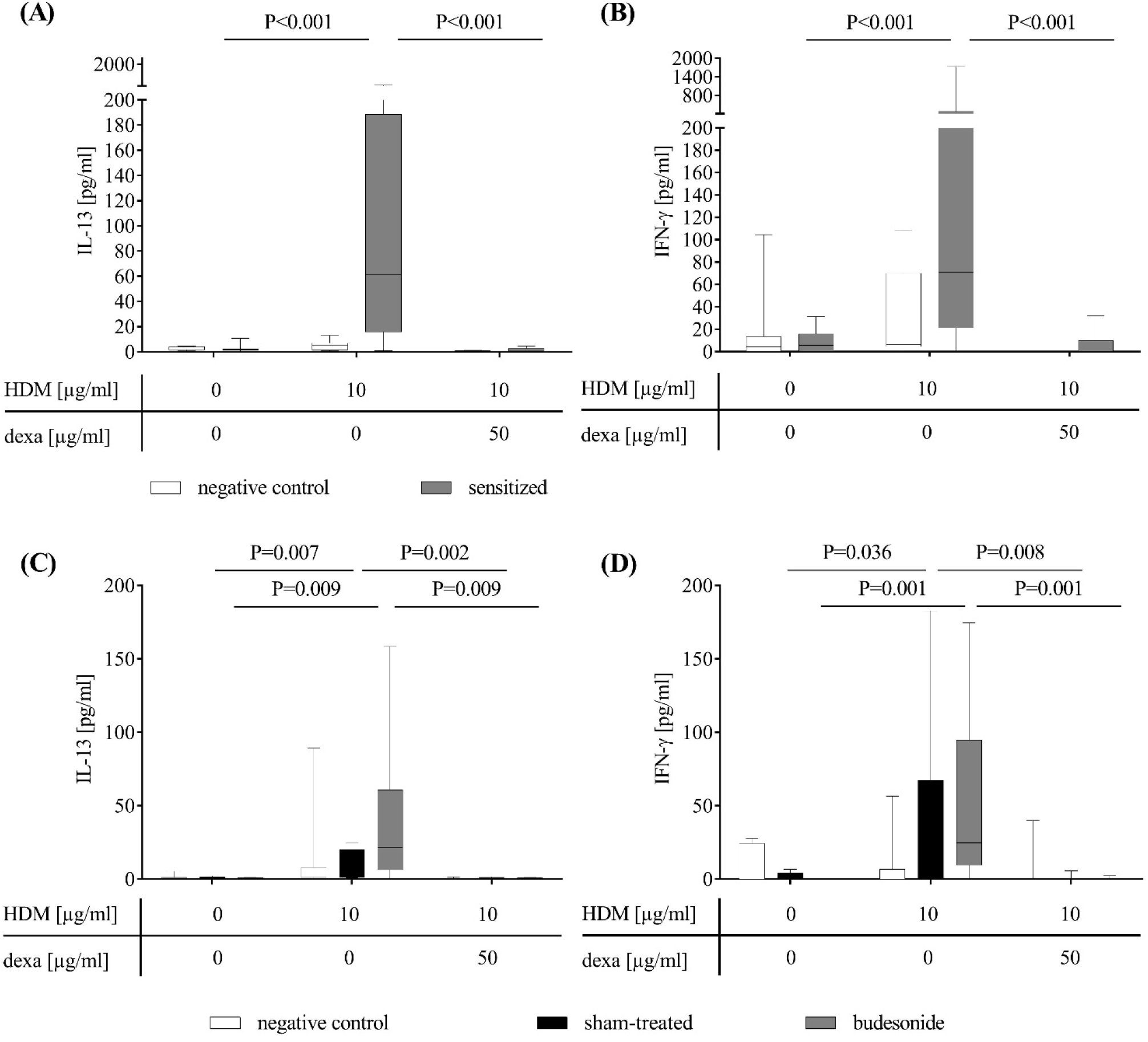
Cytokine release of restimulated PBMC after challenge and treatment. Cytokine release of HDM stimulated PBMC is depicted after challenge (A, B), and after treatment (C, D). IL-13 (A, C) and IFN-γ (B, D) were detected by ELISA after 96 h. Additional incubation with dexamethasone reduced the cytokine release independently of group membership. Box plot with median; n varies for different stimulations; HDM=house dust mite; dexa=dexamethasone; Mann-Whitney-Wilcoxon test.

**Figure S2:**
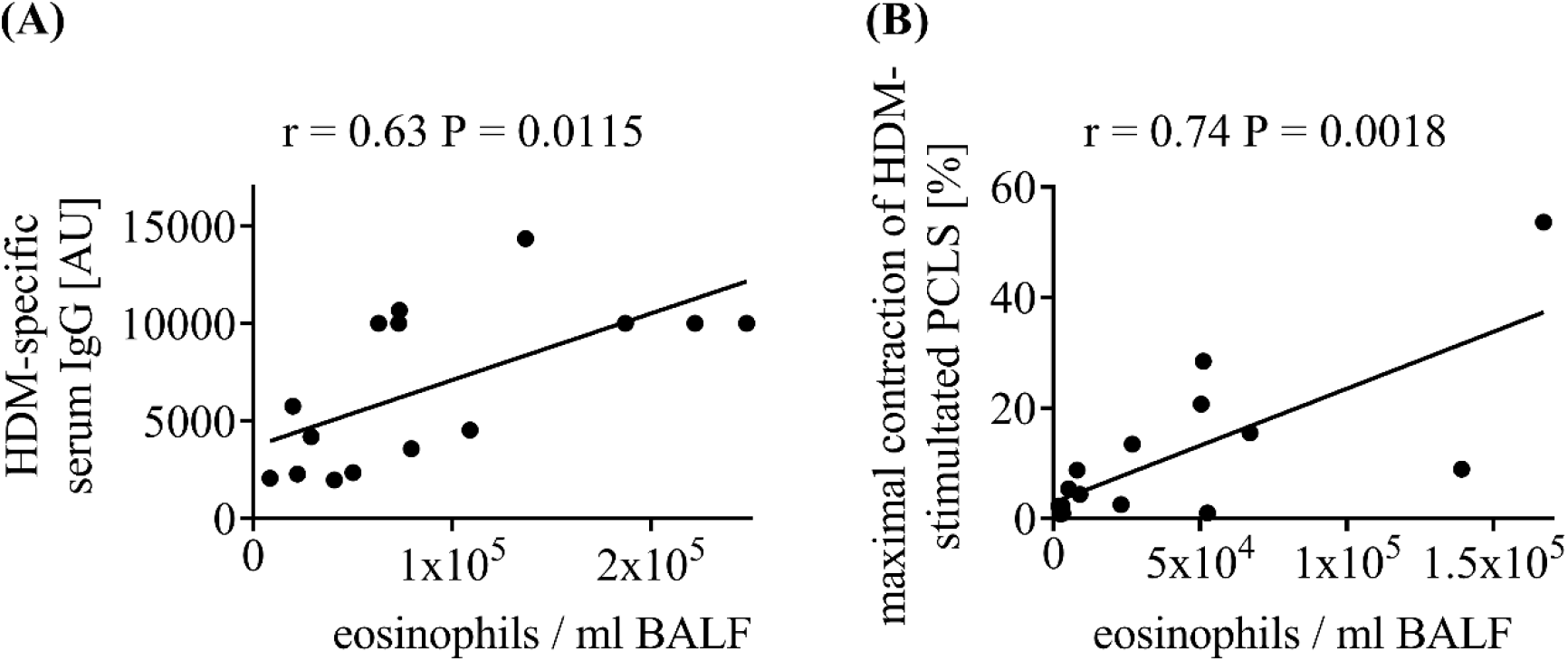
Correlation analyses for eosinophils and two other immunity-related readouts. Correlations were calculated after challenge (A) and after the therapeutic intervention (B). Eosinophils per ml BALF and HDM-specific serum IgG correlated for sensitized animals (A, n=17). After the therapeutic intervention, eosinophils per ml BALF correlated with HDM-induced PCLS-bronchoconstriction (B, n=25/15). Single values and linear regression line with corresponding Pearson correlation coefficients (r) and P values are indicated. BALF=bronchoalveolar lavage fluid; HDM=house dust mite; AU=arbitrary units; PCLS= precision-cut lung slices.

**Fig. S3:**
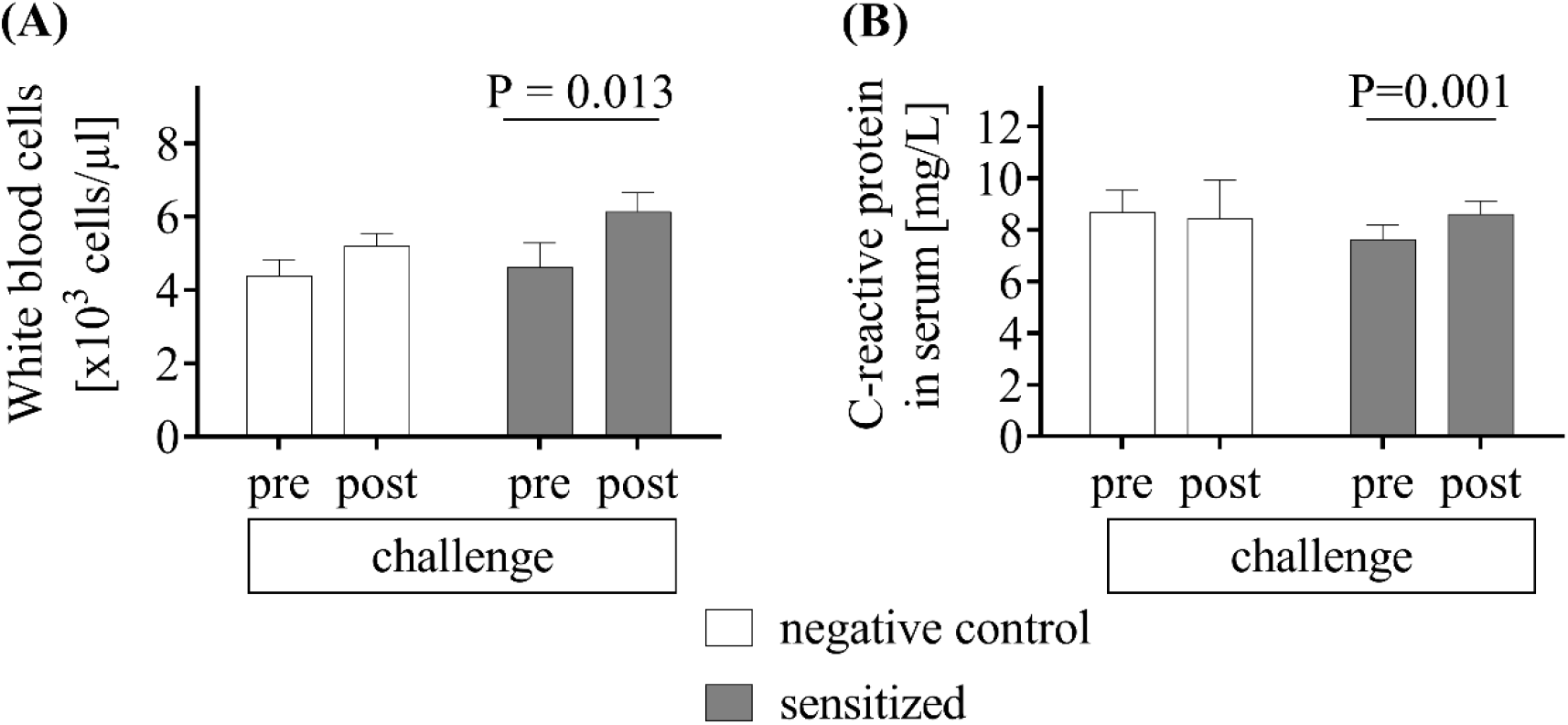
Systemic inflammatory parameters. (A) Hematological analysis of EDTA-blood reveals an increase of white blood cells only in sensitized animals (grey) after challenge compared to pre-challenge values. Mean+SEM; n=8 negative control, n=10/16 sensitized; paired t-test) (B) C-reactive protein in serum before and after HDM challenge. After the allergen challenge phase, CRP concentrations in serum were not altered in negative control animals compared to baseline. Respective levels for sensitized animals increased. Mean+SEM; n=8 negative control, n=18 sensitized; paired t-test.

## REFERENCES

1. World Health Organization (WHO). Asthma Fact sheet N°307 [Internet]. 2013 [accessed 7 March 2007]. Available from: http://www.who.int/mediacentre/factsheets/fs307/en/.

2. Brusselle G, Bracke K. Targeting immune pathways for therapy in asthma and chronic obstructive pulmonary disease. Annals of the American Thoracic Society. 2014;11 Suppl 5:S322–8. Epub 2014/12/20. doi: 10.1513/AnnalsATS.201403-118AW. PubMed PMID: 25525740.

3. Global Initiative for Asthma. Global Strategy for Asthma Management and Prevention [Report]. 2016. Available from: www.ginasthma.org.

4. Howarth PH, Bradding P, Montefort S, Peroni D, Djukanovic R, Carroll MP, et al. Mucosal inflammation and asthma. American journal of respiratory and critical care medicine. 1994;150(5 Pt 2):S18–22. Epub 1994/11/01. doi: 10.1164/ajrccm/150.5_Pt_2.S18. PubMed PMID: 7952584.

5. Zissler UM, Esser-von Bieren J, Jakwerth CA, Chaker AM, Schmidt-Weber CB. Current and future biomarkers in allergic asthma. Allergy. 2016;71(4):475–94. Epub 2015/12/27. doi: 10.1111/all.12828. PubMed PMID: 26706728.

6. Diamant Z, Gauvreau GM, Cockcroft DW, Boulet LP, Sterk PJ, de Jongh FH, et al. Inhaled allergen bronchoprovocation tests. The Journal of allergy and clinical immunology. 2013;132(5):1045–55.e6. Epub 2013/10/15. doi: 10.1016/j.jaci.2013.08.023. PubMed PMID: 24119772.

7. Wohlsen A, Martin C, Vollmer E, Branscheid D, Magnussen H, Becker WM, et al. The early allergic response in small airways of human precision-cut lung slices. The European respiratory journal. 2003;21(6):1024–32. Epub 2003/06/12. PubMed PMID: 12797499.

8. Fahy JV. Type 2 inflammation in asthma--present in most, absent in many. Nature reviews Immunology. 2015;15(1):57–65. Epub 2014/12/24. doi: 10.1038/nri3786. PubMed PMID: 25534623; PubMed Central PMCID: PMCPMC4390063.

9. Schelegle ES, Gershwin LJ, Miller LA, Fanucchi MV, Van Winkle LS, Gerriets JP, et al. Allergic asthma induced in rhesus monkeys by house dust mite (Dermatophagoides farinae). The American journal of pathology. 2001;158(1):333–41. Epub 2001/01/06. doi: 10.1016/s0002-9440(10)63973-9. PubMed PMID: 11141508; PubMed Central PMCID: PMCPMC1850255.

10. Van Scott MR, Hooker JL, Ehrmann D, Shibata Y, Kukoly C, Salleng K, et al. Dust mite-induced asthma in cynomolgus monkeys. Journal of applied physiology (Bethesda, Md : 1985). 2004;96(4):1433–44. Epub 2003/12/16. doi: 10.1152/japplphysiol.01128.2003. PubMed PMID: 14672959.

11. Iwashita K, Kawasaki H, Sawada M, In M, Mataki Y, Kuwabara T. Shortening of the Induction Period of Allergic Asthma in Cynomolgus Monkeys by Ascaris suum and House Dust Mite. Journal of Pharmacological Sciences. 2008;106(1):92–9. doi: 10.1254/jphs.FP0071523.

12. Miller LA, Plopper CG, Hyde DM, Gerriets JE, Pieczarka EM, Tyler NK, et al. Immune and airway effects of house dust mite aeroallergen exposures during postnatal development of the infant rhesus monkey. Clinical and experimental allergy : journal of the British Society for Allergy and Clinical Immunology. 2003;33(12):1686–94. Epub 2003/12/06. PubMed PMID: 14656356.

13. Ayanoglu G, Desai B, Fick RB, Jr., Grein J, de Waal Malefyt R, Mattson J, et al. Modelling asthma in macaques: longitudinal changes in cellular and molecular markers. The European respiratory journal. 2011;37(3):541–52. Epub 2010/07/24. doi: 10.1183/09031936.00047410. PubMed PMID: 20650997.

14. Van Scott MR, Reece SP, Olmstead S, Wardle R, Rosenbaum MD. Effects of acute psychosocial stress in a nonhuman primate model of allergic asthma. Journal of the American Association for Laboratory Animal Science : JAALAS. 2013;52(2):157–64. Epub 2013/04/09. PubMed PMID: 23562098; PubMed Central PMCID: PMCPMC3624783.

15. Madwed JB, Jackson AC. Determination of airway and tissue resistances after antigen and methacholine in nonhuman primates. Journal of applied physiology (Bethesda, Md : 1985). 1997;83(5):1690–6. Epub 1998/01/07. PubMed PMID: 9375340.

16. Seehase S, Schleputz M, Switalla S, Matz-Rensing K, Kaup FJ, Zoller M, et al. Bronchoconstriction in nonhuman primates: a species comparison. Journal of applied physiology (Bethesda, Md : 1985). 2011;111(3):791–8. Epub 2011/06/28. doi: 10.1152/japplphysiol.00162.2011. PubMed PMID: 21700889.

17. Joad JP, Kott KS, Bric JM, Schelegle ES, Gershwin LJ, Plopper CG, et al. The effects of inhaled corticosteroids on intrinsic responsiveness and histology of airways from infant monkeys exposed to house dust mite allergen and ozone. Toxicology and applied pharmacology. 2008;226(2):153–60. Epub 2007/11/10. doi: 10.1016/j.taap.2007.09.005. PubMed PMID: 17991502.

18. Curths C, Knauf S, Kaup F-J. Respiratory Animal Models in the Common Marmoset (Callithrix jacchus). Veterinary Sciences. 2014;1(1):63–76. doi: 10.3390/vetsci1010063.

19. Seehase S, Lauenstein HD, Schlumbohm C, Switalla S, Neuhaus V, Forster C, et al. LPS-induced lung inflammation in marmoset monkeys - an acute model for anti-inflammatory drug testing. PloS one. 2012;7(8):e43709. doi: 10.1371/journal.pone.0043709. PubMed PMID: 22952743; PubMed Central PMCID: PMC3429492.

20. Curths C, Wichmann J, Dunker S, Windt H, Hoymann HG, Lauenstein HD, et al. Airway hyper-responsiveness in lipopolysaccharide-challenged common marmosets (Callithrix jacchus). Clinical science. 2014;126(2):155–62. doi: 10.1042/CS20130101. PubMed PMID: 23879175; PubMed Central PMCID: PMC3793853.

21. Rondeau JM, Ramage P, Zurini M, Gram H. The molecular mode of action and species specificity of canakinumab, a human monoclonal antibody neutralizing IL-1beta. mAbs. 2015;7(6):1151–60. Epub 2015/08/19. doi: 10.1080/19420862.2015.1081323. PubMed PMID: 26284424; PubMed Central PMCID: PMCPMC4966334.

22. European Medicines Agency (EMeA). CHMP assessment report for ILARIS. International Nonproprietary Name: canakinumab. Procedure No. EMEA/H/C/001109 2009 EMEA/503722/2009. Available from: http://www.ema.europa.eu/docs/en_GB/document_library/EPAR_-_Public_assessment_report/human/001109/WC500031679.pdf.

23. Fuchs B, Braun A. Improved mouse models of allergy and allergic asthma--chances beyond ovalbumin. Current drug targets. 2008;9(6):495–502. Epub 2008/06/10. PubMed PMID: 18537588.

24. Nials AT, Uddin S. Mouse models of allergic asthma: acute and chronic allergen challenge. Disease models & mechanisms. 2008;1(4-5):213–20. Epub 2008/12/19. doi: 10.1242/dmm.000323. PubMed PMID: 19093027; PubMed Central PMCID: PMCPMC2590830.

25. Yasue M, Nakamura S, Yokota T, Okudaira H, Okumura Y. Experimental monkey model sensitized with mite antigen. International archives of allergy and immunology. 1998;115(4):303–11. Epub 1998/05/05. PubMed PMID: 9566353.

26. Lommatzsch M, Julius P, Kuepper M, Garn H, Bratke K, Irmscher S, et al. The course of allergen-induced leukocyte infiltration in human and experimental asthma. The Journal of allergy and clinical immunology. 2006;118(1):91–7. Epub 2006/07/04. doi: 10.1016/j.jaci.2006.02.034. PubMed PMID: 16815143.

27. Ray A, Kolls JK. Neutrophilic Inflammation in Asthma and Association with Disease Severity. Trends Immunol. 2017;38(12):942–54. Epub 2017/08/09. doi: 10.1016/j.it.2017.07.003. PubMed PMID: 28784414; PubMed Central PMCID: PMCPMC5711587.

28. Young SS, Ritacco G, Skeans S, Chapman RW. Eotaxin and nitric oxide production as markers of inflammation in allergic cynomolgus monkeys. International archives of allergy and immunology. 1999;120(3):209–17. Epub 1999/12/11. doi: 24269. PubMed PMID: 10592466.

29. Schelegle ES, Miller LA, Gershwin LJ, Fanucchi MV, Van Winkle LS, Gerriets JE, et al. Repeated episodes of ozone inhalation amplifies the effects of allergen sensitization and inhalation on airway immune and structural development in Rhesus monkeys. Toxicology and applied pharmacology. 2003;191(1):74–85. Epub 2003/08/14. PubMed PMID: 12915105.

30. Wang L, Jenkins TJ, Dai M, Yin W, Pulido JC, Lamantia-Martin E, et al. Antagonism of chemokine receptor CCR8 is ineffective in a primate model of asthma. Thorax. 2013;68(6):506–12. Epub 2013/03/05. doi: 10.1136/thoraxjnl-2012-203012. PubMed PMID: 23457038.

31. Laan MP, Baert MR, Vredendaal AE, Savelkoul HF. Differential mRNA expression and production of interleukin-4 and interferon-gamma in stimulated peripheral blood mononuclear cells of house-dust mite-allergic patients. European cytokine network. 1998;9(1):75–84. Epub 1998/06/05. PubMed PMID: 9613681.

32. Jenmalm MC, Bjorksten B. Development of immunoglobulin G subclass antibodies to ovalbumin, birch and cat during the first eight years of life in atopic and non-atopic children. Pediatric allergy and immunology : official publication of the European Society of Pediatric Allergy and Immunology. 1999;10(2):112–21. Epub 1999/09/09. PubMed PMID: 10478613.

33. Miranda DO, Silva DA, Fernandes JF, Queiros MG, Chiba HF, Ynoue LH, et al. Serum and salivary IgE, IgA, and IgG4 antibodies to Dermatophagoides pteronyssinus and its major allergens, Der p1 and Der p2, in allergic and nonallergic children. Clinical & developmental immunology. 2011;2011:302739. Epub 2011/10/19. doi: 10.1155/2011/302739. PubMed PMID: 22007250; PubMed Central PMCID: PMCPMC3189464.

34. Shimoda T, Obase Y, Kishikawa R, Iwanaga T. Serum high-sensitivity C-reactive protein can be an airway inflammation predictor in bronchial asthma. Allergy and asthma proceedings. 2015;36(2):e23–8. Epub 2015/02/26. doi: 10.2500/aap.2015.36.3816. PubMed PMID: 25715235.

35. Wang X, Reece S, Olmstead S, Wardle RL, Van Scott MR. Nocturnal thoracoabdominal asynchrony in house dust mite-sensitive nonhuman primates. Journal of asthma and allergy. 2010;3:75–86. Epub 2010/01/01. PubMed PMID: 21437042; PubMed Central PMCID: PMCPMC3047915.

36. Hovenberg HW, Davies JR, Carlstedt I. Different mucins are produced by the surface epithelium and the submucosa in human trachea: identification of MUC5AC as a major mucin from the goblet cells. The Biochemical journal. 1996;318 (Pt 1):319–24. Epub 1996/08/15. PubMed PMID: 8761488; PubMed Central PMCID: PMCPMC1217624.

37. Seidel V. Morphological investigations on the lung of common marmosets (Callithrix jacchus) (in German). [Doctoral Thesis]: Doctoral Thesis, School of Veterinary Medicine, Hannover, Germany; 2012.

38. Roth FD, Quintar AA, Uribe Echevarria EM, Torres AI, Aoki A, Maldonado CA. Budesonide effects on Clara cell under normal and allergic inflammatory condition. Histochem Cell Biol. 2007;127(1):55–68. Epub 2006/07/22. doi: 10.1007/s00418-006-0220-3. PubMed PMID: 16858555.

39. Shijubo N, Itoh Y, Yamaguchi T, Imada A, Hirasawa M, Yamada T, et al. Clara cell protein-positive epithelial cells are reduced in small airways of asthmatics. American journal of respiratory and critical care medicine. 1999;160(3):930–3. Epub 1999/09/03. doi: 10.1164/ajrccm.160.3.9803113. PubMed PMID: 10471621.

